# Partially hydrolyzed guar gum attenuates the symptoms of SARS-CoV-2 infection through gut microbiota modulation in an animal model

**DOI:** 10.1101/2023.06.13.544519

**Authors:** Jiayue Yang, Isaiah Song, Misa Saito, Tenagy Hartanto, Takeshi Ichinohe, Shinji Fukuda

## Abstract

The coronavirus disease 2019 (COVID-19) pandemic has caused worldwide health issues. Although several vaccines have been developed, it is still difficult to prevent and reduce the inflammation caused by the infection. Studies have shown that there are correlations between the gut environment and severity of symptoms caused by SARS-CoV-2 infection. Several gut metabolites produced by the gut microbiota such as SCFAs and the secondary bile acid UDCA are reported to improve the survival rate of the host after viral infection in an animal model through modulation of the host immune system. Therefore, in this study, we attempted to use the prebiotic dietary fiber PHGG to modulate the gut microbiome and intestinal metabolites for improvement of host survival rate after SARS-CoV-2 infection in a Syrian hamster model. We were able to show that PHGG significantly improved the host survival rate and body weight reduction. Analysis of the gut microbiome, serum, and intestinal metabolites revealed that PHGG significantly increased the concentrations of several intestinal SCFAs, fecal secondary bile acids, and serum secondary bile acids. Furthermore, several microbial species and metabolites identified in this study are consistent with reports in humans. Taken together, our data suggest that PHGG is a candidate prebiotic food for reducing the morbidity of COVID-19.

## Introduction

Coronavirus disease 2019 (COVID-19) pandemic has caused major health issues in society. Severe acute respiratory syndrome coronavirus 2 (SARS-CoV-2) infection-induced cytokine storms can lead to severe morbidity and mortality ^1^. Although there are several factors reported to be correlated with severity of symptoms ^2^. The severity of symptoms are still unpredictable, and treatment options are limited. Numerous people suffer from the long-term effects of the infection ^3^. Despite the development of vaccines, new SARS-CoV-2 variants continue to emerge and escape the adaptive immune response elicited by vaccination ^4^. Therefore, it is necessary to find ways to prevent and reduce the inflammation and cytokine storm caused by the infection.

It has been reported that the gut microbiota is important for health. They produce many kinds of metabolites such as short-chain fatty acids (SCFAs) and secondary bile acids, which have various biological functions such as host immune system modulation and anti-inflammation ^5-7^. Studies have shown that there is crosstalk between the gut microbiota and pulmonary system, which is referred to as the gut-lung axis ^8^. In particular, there are several reports of a link between the gut microbiota and SARS-CoV-2 infection. The gut microbiome of patients with COVID-19 show low diversity and patients with COVID-19 possess fewer SCFA producers ^9,10^. Recently, the SCFA acetic acid has been recognized for its ability to inactivate SARS-CoV-2 in an *in vitro* experiment ^11^. It has been reported that the gut microbiota-derived secondary bile acid ursodeoxycholic acid (UDCA) is able to reduce the expression of viral host receptor angiotensin-converting enzyme 2 (ACE2) through protein-ligand interaction with farnesoid X receptors (FXR) ^4^. Furthermore, UDCA treatment has been correlated with positive clinical outcomes in COVID-19 cases ^4^.

Gut microbiome profiles are influenced by the daily diet of the host ^12^. The correlation between diet and COVID-19 has been reported in the literature ^1^. It is well-known that the gut microbiota produces SCFAs from dietary fiber ^7^. Vegetarian patients have been associated with decreased severity of COVID-19-related inflammation ^13^. In addition, one large prospective survey has revealed that a plant-rich diet reduces the risk and severity of COVID-19 ^14^. Therefore, we considered that dietary intervention may contribute to the prevention of inflammation caused by SARS-CoV-2 infection. In this study, we attempted to use the prebiotic dietary fiber partially hydrolyzed guar gum (PHGG) to modulate the gut microbiome and its metabolites for the prevention and attenuation of the inflammation caused by SARS-CoV-2 infection. PHGG is a water-soluble, low-viscosity dietary fiber ^15^. It is manufactured by enzymatic hydrolysis of high viscose dietary fiber guar gum that is made from *Cyamopsis tetragonoloba*, also called cluster beans or guar, which grows in tropical and subtropical regions and is widely cultivated in India, Pakistan, and the USA ^16^. It is used as a prebiotic food for the improvement of constipation ^17^. A previous study reported that PHGG is able to alter the gut microbiome and metabolite compositions by increasing the abundance of functional bacteria such as *Bifidobacterium* as well as intestinal SCFA concentrations ^18^. Moreover, in a human clinical study, PHGG has been reported to prevent influenza infection ^19^. Therefore, we considered that PHGG may be able to increase resistance to SARS-CoV-2 infection. We used a SARS-CoV-2 hamster infection experimental model to examine the potential preventative effects of PHGG. We also analyzed the changes in the gut microbiome and gut metabolites as a result of PHGG consumption. Consequently, our data showed that PHGG significantly reduced mortality and improved recovery after SARS-CoV-2 infection. Therefore, we propose that PHGG be further investigated as a prebiotic that can potentially reduce the inflammation caused by SARS-CoV-2 infection.

## Results

### PHGG diet attenuated SARS-CoV-2 infection

SARS-CoV-2 infects humans through ACE2 receptors expressed in the lungs, blood vessels, and intestinal epithelia ^11^. Hamsters possess ACE2 receptors and are commonly used for SARS-CoV-2 infection animal experiments ^20^. This model is able to recapitulate several hallmark features of COVID-19 in humans such as inflammation in the lungs and changes in the gut microbiota and gut metabolites ^21^. Therefore, in this study, we used Syrian hamsters to assess the preventative effects of PHGG on SARS-CoV-2 infection. Hamsters were given a PHGG-supplemented diet or control diet for two weeks before SARS-CoV-2 infection (Fig. 1A). Then, each hamster was exposed to SARS-CoV-2 (Fig. 1A). As a result, all hamsters in the PHGG group survived after exposure to SARS-CoV-2 while the control group showed a poor survival rate (Fig. 1B). Moreover, compared to the control group, the PHGG group more rapidly recovered their body weight (Fig. 1C). These results indicate that PHGG can reduce morbidity and promote recovery from the inflammation caused by SARS-CoV-2 infection.

**Figure 1.**
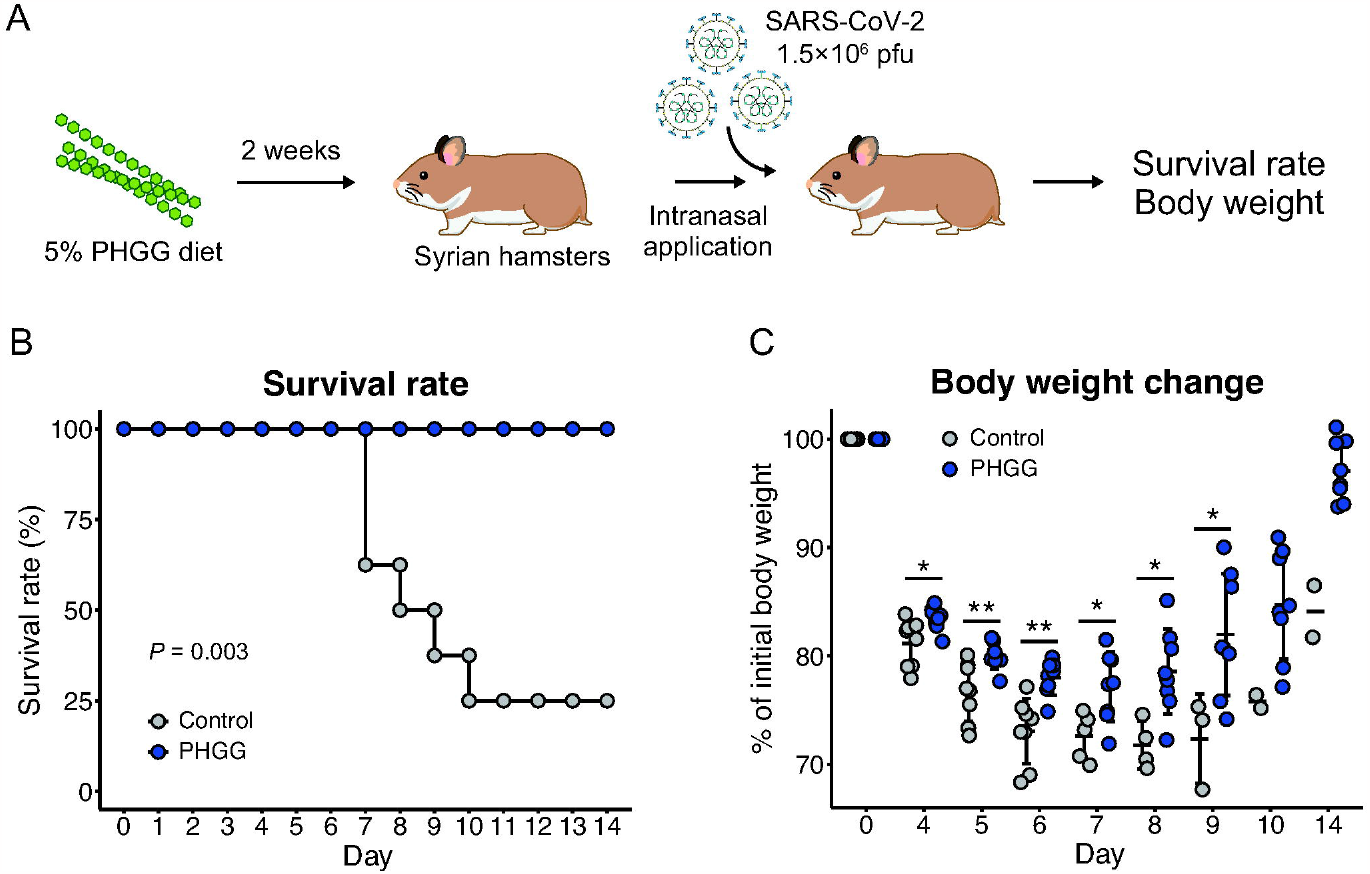
PHGG diet attenuated the survival rate and body weight reduction caused by SARS-CoV-2 infection in Syrian hamsters. (A) Experimental procedures of SARS-CoV-2 infection experiment using Syrian hamsters are shown. (B) Survival rates of hamsters after exposure to SARS-CoV-2. **, *P* < 0.01 (generalized Wilcoxon test) (C) Body weight change of hamsters after exposure to SARS-CoV-2. *, *P* < 0.05 **, *P* < 0.01 (Wilcoxon rank sum test).

### PHGG diet altered the gut microbiome profile

To elucidate the mechanism of this inflammation alleviatory effect by PHGG, we performed microbiome analysis of the cecal contents by 16S rRNA gene amplicon sequencing using next-generation sequencing. UniFrac analysis and analysis of similarity (ANOSIM) showed that the PHGG diet significantly altered the gut microbiome profile of hamsters (Fig. 2AB). The relative abundance of *Ileibacterium, Bifidobacterium*, and *Prevotella* genera were significantly increased in the PHGG group, while *Alistipes* and *Desulfovibrio* were significantly decreased (Fig. 2CD, Fig. 3). This data indicates that the PHGG diet altered the gut microbiome profile of hamsters.

**Figure 2.**
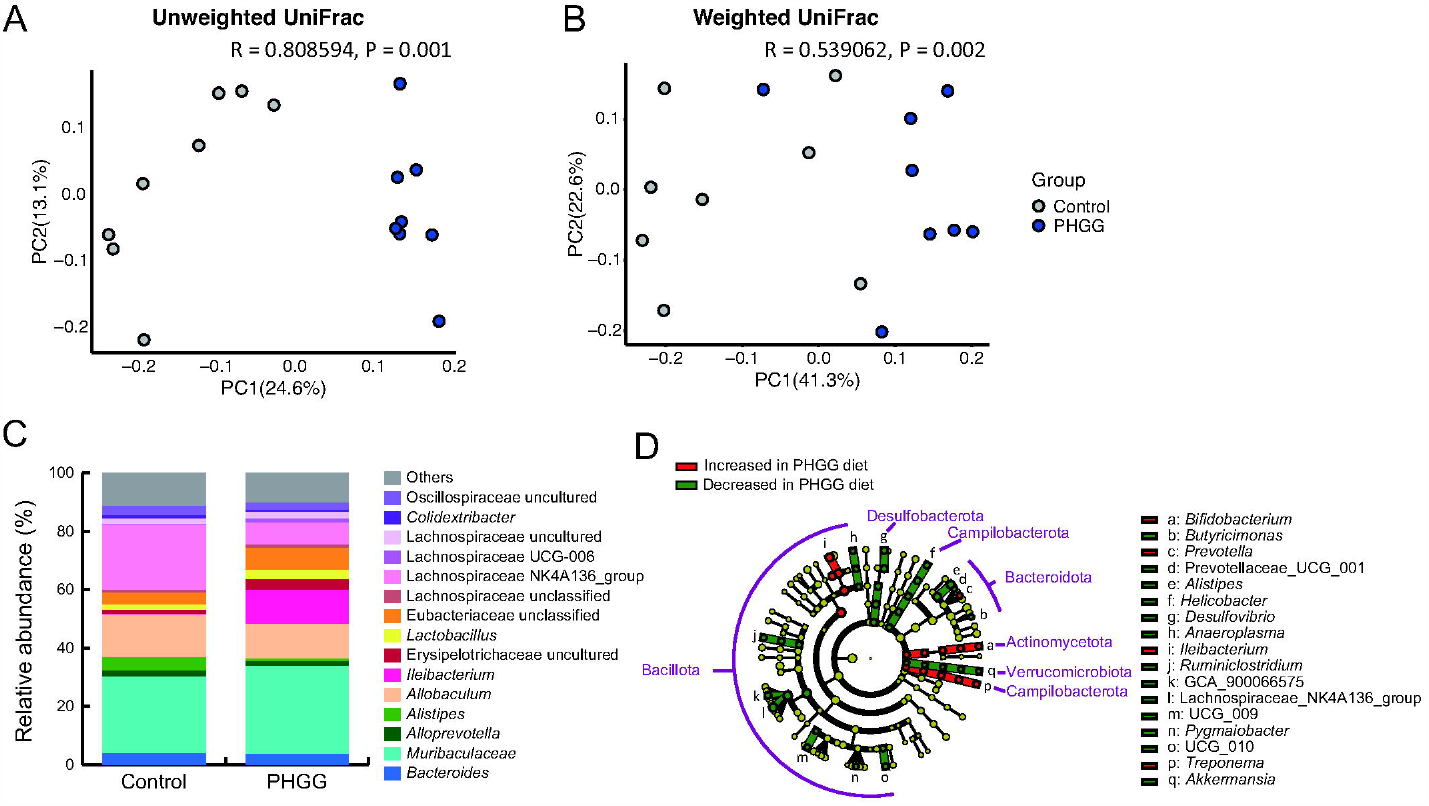
PHGG diet altered the gut microbiome profile of Syrian hamsters. (A) Unweighted and (B) weighted UniFrac analyses and ANOSIM of the gut microbiome profile in both control and PHGG groups. (C) Relative abundance of the gut microbiome. (D) LEfSe analysis of the gut microbiome profile.

**Figure 3.**
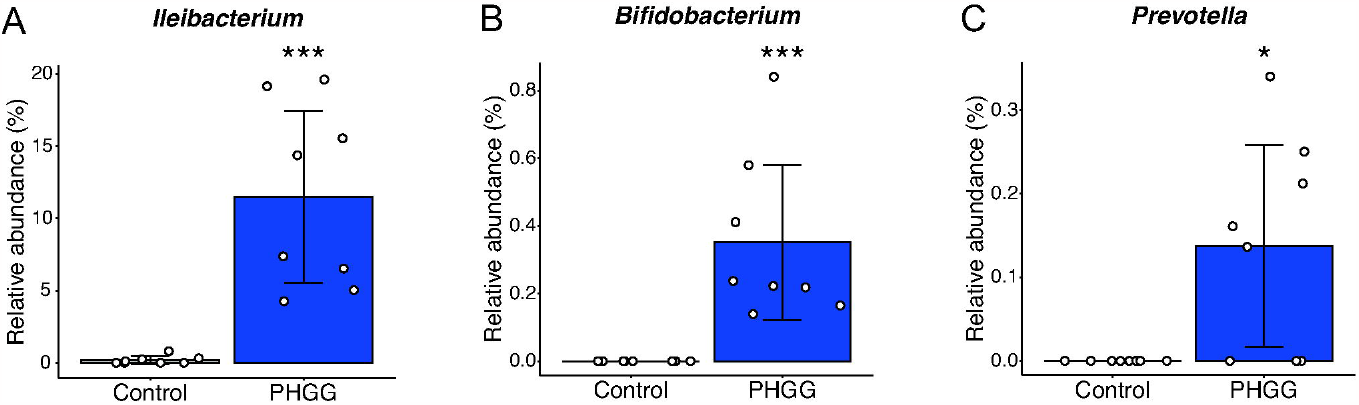
SCFA producers increased in PHGG group. Relative abundance of (A) *Ileibacterium*, (B) *Bifidobacterium*, and (C) *Prevotella*. *, *P* < 0.05 ***, *P* < 0.001 (Wilcoxon rank sum test).

### PHGG diet altered the intestinal metabolome profile

Since *Bifidobacterium* and *Prevotella* are known SCFA producers, we performed metabolome analysis by gas chromatography mass spectrometry (GC/MS) and liquid chromatography mass spectrometry (LC-MS) to observe SCFA and bile acid production in the gut environment. According to the results, the total amount of SCFAs was significantly increased in the PHGG group (Fig. 4A, Table S1). Among the SCFAs, propionic acid and valeric acid were significantly increased in the PHGG group (Fig. 4BC, Table S1), while formic acid was decreased (Fig. 4D, Table S1). We analyzed the primary and secondary bile acid profile in the feces using LC-MS as well, as they are reported to have anti-inflammatory effects and the potential to attenuate the severity of COVID-19 ^7,22^. As a result, the amounts of anti-inflammatory secondary bile acids UDCA and DCA were found to be increased in the feces and serum of PHGG group hamsters, respectively (Fig. 4E, Table S2). In particular, DCA, which has been reported to have anti-infective effects against SARS-CoV-2 ^23^, was significantly higher in the serum of PHGG group hamsters (Fig. 4F, Table S2).

**Figure 4.**
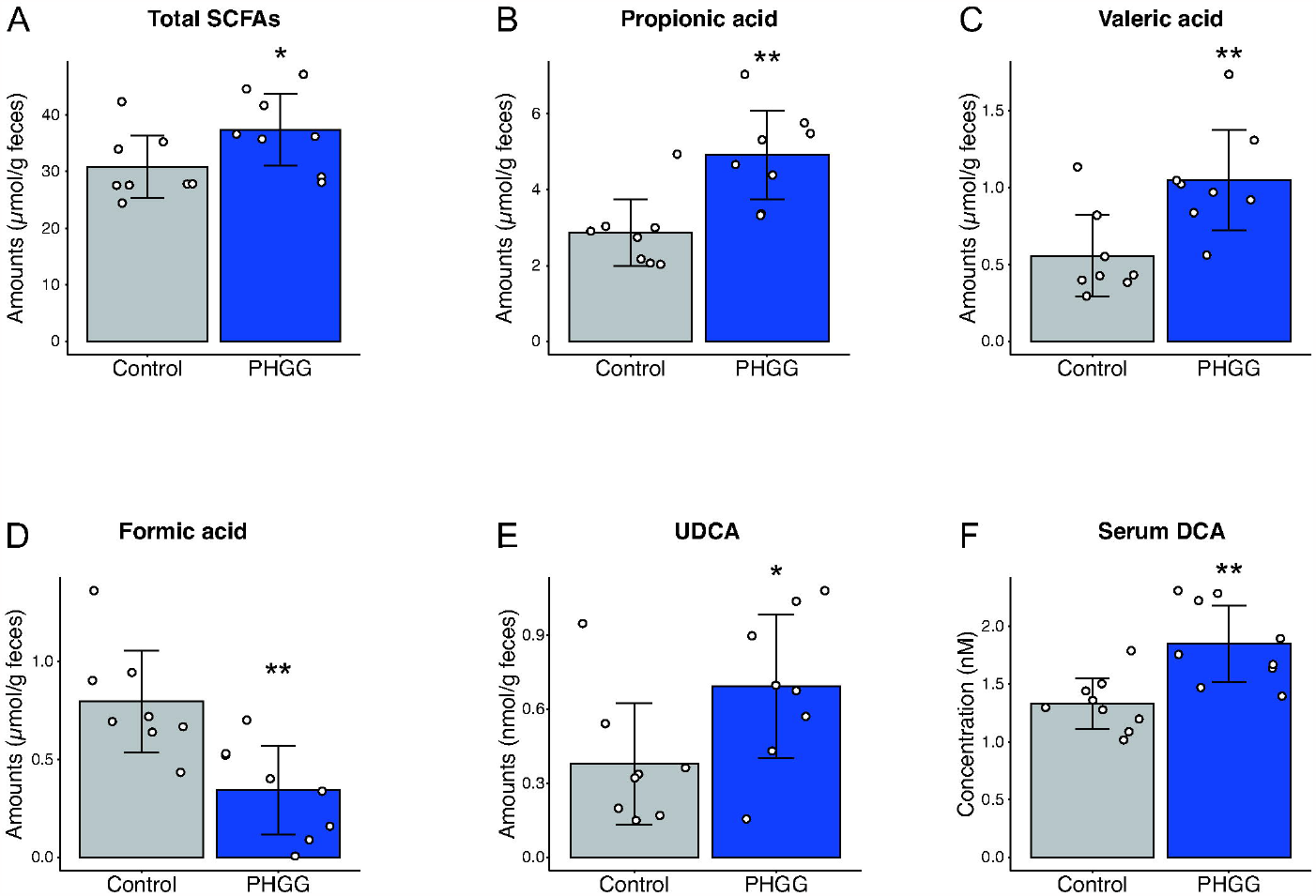
PHGG diet altered SCFAs and secondary bile acids profiles in the feces of hamsters. Comparisons of SCFA concentrations in feces altered in PHGG group: (A) Total SCFAs (B) Propionate (C) Valerate (D) Formate. (E) Comparison of UDCA in feces. (F) Comparison of DCA in serum. *, *P* < 0.05 **, *P* < 0.01 (Wilcoxon rank sum test).

In order to elucidate the changes in intestinal metabolome profile associated with the specific members of the gut microbiota that were found to be altered in the PHGG group, we performed correlation analysis of *Ileibacterium, Bifidobacterium*, and *Prevotella* against SCFAs and secondary bile acids in feces and serum. We found that *Ileibacterium* was positively correlation with valeric acid, while it was negative correlated with formic acid (Fig. 5AB). Interestingly, *Bifidobacterium* showed a positive correlation with propionic acid, although propionic acid production ability by *Bifidobacterium* has not been reported in the literature (Fig. 5C).

**Figure 5.**
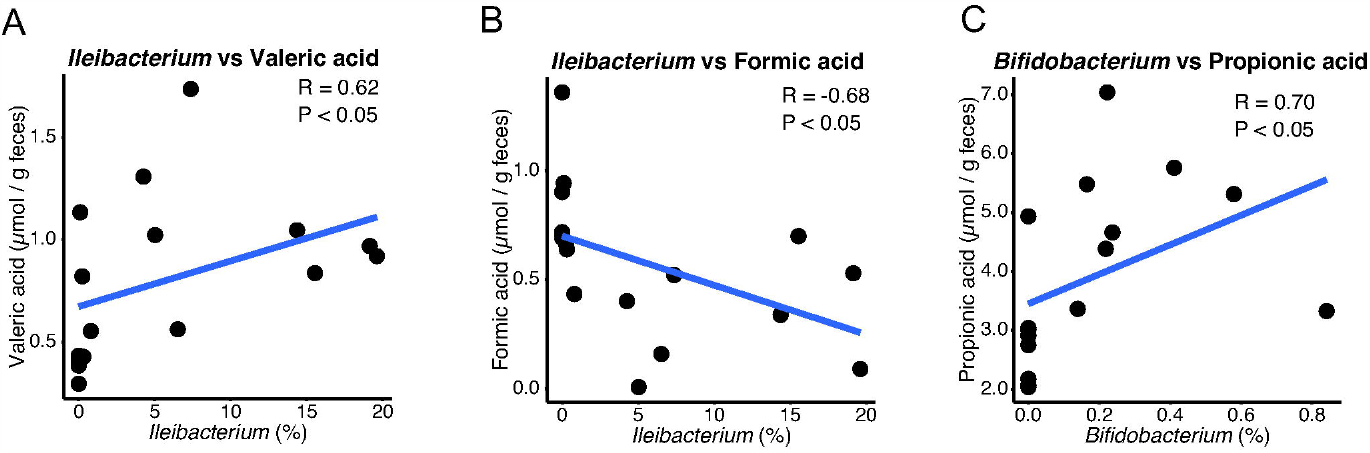
Gut microbes correlated with the amount of SCFAs. Correlation analysis between *Ileibacterium* and (A) valerate or (B) formate, (C) *Bifidobacterium* with propionate.

## Discussion

Our data indicates that PHGG administration significantly suppressed morbidity and mortality in SARS-CoV-2 infected Syrian hamsters. Our data also showed that PHGG administration altered the gut microbiome profile and gut metabolites of Syrian hamsters. Studies have suggested that dietary intervention may increase resistance to SARS-CoV-2 infection through modulation of the gut microbiota and metabolites ^24-26^. Our data further supports that notion by showing that modulation of the gut microbiota by dietary fiber contributes to positive outcomes in response to SARS-CoV-2 infection.

In our data, relative abundances of *Ileibacterium, Bifidobacterium*, and *Prevotella* were significantly increased in the PHGG diet group (Fig. 3). *Ileibacterium* has been reported to be associated with polysaccharide and SCFAs metabolism and is believed to be a producer of SCFAs ^27-29^. Thus, it is possible that PHGG increases the abundance of *Ileibacterium*, resulting in a higher amount of SCFAs in the intestines. As a matter of fact, the total amount of SCFAs was increased in the PHGG diet group. In several human clinical studies, the relative abundance of *Bifidobacterium* was also increased in those consuming a PHGG-supplemented diet ^18,30^. A PHGG diet also increased SCFA concentrations in hamsters (Fig. 4A), similarly to human study results ^18^. Valeric acid and propionic acid, which were increased in the PHGG group, are known for their anti-inflammation effects ^31,32^. In our study, valeric acid was positively correlated with *Ileibacterium* (Fig. 5A). Although *Ileibacterium* has been reported for its association with SCFA metabolism, it has not yet been reported to produce valeric acid. Therefore, further investigation is necessary in the future. Propionic acid was positively correlated with *Bifidobacterium*. It is conceivable that the cross-feeding of *Bifidobacterium* with other bacteria may contribute to the increase of production of this SCFA, as *Bifidobacterium* has not been reported to produce propionic acid. On the other hand, formic acid, which was decreased in the PHGG diet group, has been reported to be elevated in inflammation-associated dysbiosis ^33^.

Gut microbes convert primary bile acids to various secondary bile acids ^34^. The PHGG diet increased the amount of the secondary bile acid UDCA in feces. A previous study showed that UDCA reduces ACE2 expression and is correlated with positive clinical outcomes in COVID-19 treatment ^4^. Additionally, DCA was increased in the serum of the PHGG diet group. As mentioned before, DCA has been reported to have anti-infective effects against SARS-CoV-2 ^23^. As both UDCA and DCA are ligands of FXR ^4,35^, we hypothesize that PHGG consumption enhances resistance to SARS-CoV-2 infection through FXR binding, as activation of this receptor is found to reduce the expression of ACE2 ^4^.

Several studies have suggested that probiotic bacteria such as *Bifidobacterium* and *Lactobacillus* may improve the clinical outcome of SARS-CoV-2 infection ^26,36^. A clinical trial using encapsulated synbiotic formula SIM01 consisting of 3 *Bifidobacterium* strains and 3 prebiotic polysaccharides suggested that SIM01 has the potential to increase resistance to SARS-CoV-2 infection ^24^. Compared to probiotic bacteria, the production cost of prebiotic dietary fiber PHGG is likely lower, as well as being easier to transport and consume. Thus, PHGG may be more accessible for use in daily life but further studies are first needed to clarify the effects of PHGG in human clinical outcomes.

Nevertheless, our study suggests that PHGG has great potential to improve the symptoms of SARS-CoV-2 infection. Despite the development of COVID-19 vaccines, SARS-CoV-2 is still widespread and continues to cause problems. We speculate that dietary consumption of PHGG presents a simple, easily accessible way to help resist infection.

## Materials and methods

### Animal experiment

Four-week-old female Syrian hamsters were purchased from Japan SLC, Inc. PHGG was purchased from Nestle Japan. Hamsters were given a 5% (w/w) PHGG-supplemented AIN-93G diet (CLEA Japan, Inc., 5% (w/w) corn starch was replaced with 5% (w/w) PHGG) or AIN-93G control diet for two weeks. After that, feces and serum of each group were collected and the microbiome profile and metabolites were analyzed. SARS-CoV-2 exposure was performed by intranasal application of viral suspension (150 μL PBS containing 1.5×10^6^ pfu of an ancestral SARS-CoV-2 strain) to hamsters under anaesthesia. Hamsters were considered to have reached the endpoint at 70% of starting weight. All experiments were performed in enhanced biosafety level 3 (BLS-3) containment laboratories at the University of Tokyo, in accordance with the institutional biosafety operating procedures.

### DNA extraction

The DNA of fecal samples was extracted according to the following steps. First, fecal samples were lyophilized by a VD-800R lyophilizer (TAITEC) for at least 18 h. Freeze-dried feces (10 mg) were suspended in 300 μl of 10% (w/v) SDS/TE (10mM Tris-HCl, 1mM EDTA, and pH 8.0) solution and 300 μl of phenol/chloroform/isoamyl alcohol (25:24:1). Then, the mixture was homogenized with 3.0 mm and 0.1 mm zirconia beads by ShakeMaster® NEO homogenizer (Biomedical Science, Tokyo, Japan) for 15 min at 1,500 × g. After that, samples were centrifuged for 10 min at max speed and 200 μl of the aqueous phase was used for DNA extraction. DNA extraction was performed by the automated DNA Extraction system GENE PREP STAR PI-480 (Kurabo Industries Ltd.) according to the manufacturer’s protocol.

### Gut microbiome analysis

Gut microbiome profile of stool samples was analyzed by 16S rRNA amplicon sequencing with the following procedure. The V1-V2 variable region of stool DNA was amplified by universal primer set 27F-mod (5’-AGRGTTTGATYMTGGCTCAG-3’) and 338R (5’-TGCTGCCTCCCGTAGGAGT-3’) using Gflex DNA polymerase (Takara) ^37^. After that, the amplified DNA products were sequenced by next-generation sequencer Miseq (Illumina). Gut microbiome profile was analyzed by Qiime2 (version 2021.11). Sequence data were trimmed and processed by using the DADA2 pipeline for quality filtering and denoising (options: –p-trunc-len-f 285 –p-trunc-len-r 215). The denoised sequences were assigned to taxa using the Silva SSU Ref Nr 99 (version 138) database with the “qiime feature-classifier classify-sklearn” command with default parameters. Unifrac distance were calculated using 12212 reads per sample with “qiime diversity core-metrics-phylogenetic” command.

### Measurement of SCFAs

Fecal samples were lyophilized by a VD-800R lyophilizer (TAITEC) for at least 18 h. Freeze-dried feces were homogenized with 3.0 mm zirconia beads by ShakeMaster® NEO homogenizer (Biomedical Science, Tokyo, Japan) for 10 min at 1,500 × g. 10 mg of fecal samples were used for the analysis. Measurement of SCFAs (formate, acetate, propionate, isobutyrate, butyrate, isovalerate, valerate, lactate, and succinate) was performed by gas chromatography mass spectrometry (GC/MS) using previously described methods ^38^.

### Measurement of bile acids

Fecal samples were lyophilized by a VD-800R lyophilizer (TAITEC) for at least 18 h. Freeze-dried feces were homogenized with 3.0 mm zirconia beads by ShakeMaster® NEO homogenizer (Biomedical Science, Tokyo, Japan) for 10 min at 1,500 × g. 10 mg of fecal sample or 50 μl of serum sample were used for the analyses. Measurement of bile acids was performed by liquid chromatography mass spectrometry (LC-MS) using previously described methods ^38^.

### Statistical analyses

Statistical analyses and correlation analysis (method: Spearman) were performed using R (version 4.2.2). Survival rate was compared using the survival package of R and generalized Wilcoxon test. The Wilcoxon-Rank-Sum test was used to compare the other data of the two groups.

## Data availability

The microbiome analysis data have been deposited in the DNA Data Bank of Japan (DDBJ) Sequence Read Archive (http://trace.ddbj.nig.ac.jp/dra/) as DRA016419.

## Acknowledgements

We thank Yoshihiro Kawaoka (University of Wisconsin and University of Tokyo) for providing SARS-CoV-2/UT-NCGM02/Human/2020/Tokyo. This work was supported in part by research grants from JSPS KAKENHI (22H03541 to S.F.), AMED-CREST (JP23gm1010009 to S.F.), JST ERATO (JPMJER1902 to S.F.), the Food Science Institute Foundation (to S.F.), and AMED (JP223fa627001 to T.I.). Fecal microbiome and metabolome analyses were funded in part by Nestle Japan.

## Conflicts of interest

S.F. is a founder and CEO and M.S. and T.H. are members of Metagen, Inc., a company involved in microbiome-based healthcare. The other authors declare no competing interests.

## Author contribution

Conceptualization: S.F.; Animal experiment: T.I.; Microbiome and metabolome analyses: M.S. and T.H.; Formal analysis: J.Y.; Writing – Original Draft: J.Y. and I.S.; Writing – Review & Editing: J.Y., I.S., T.I., and S.F.

