## Supplementary Tables for "Partially hydrolyzed guar gum attenuates the symptoms of SARS-CoV-2 infection through gut microbiota modulation in an animal model"

**Supplementary Table 1.** Amounts of SCFAs in feces.

| SCFAs | Control (nmol / g feces) Average ± Standard deviation | PHGG (nmol / g feces) Average ± Standard deviation | *P* value |
| --- | --- | --- | --- |
| Acetic acid | 22326.6 ± 3794.6 | 25358.5 ± 4016.5 | N.S. |
| Butyric acid | 3552.0 ± 3159.8 | 5069.7 ± 1896.2 | N.S. |
| Propionic acid | 2864.9 ± 876.8 | 4916.5 ± 1171.9 | ** |
| Formic acid | 794.6 ± 259.9 | 343.4 ± 226.0 | ** |
| Valeric acid | 556 ± 263.9 | 1049.8 ± 325.0 | ** |
| Succinic acid | 278.7 ± 166.7 | 386.3 ± 261.4 | N.S. |
| Lactic acid | 188.1 ± 279. 0 | 21.6 ± 20.8 | N.S. |
| Isobutyric acid | 160.8 ± 51.0 | 139.8 ± 66.9 | N.S. |
| Isovaleric acid | 97.4 ± 41.6 | 75.9 ± 34.1 | N.S. |
| Total | 30819.3 ± 5501.6 | 37361.4 ± 6389.3 | * |

Average values ± standard deviation for each SCFA and group are shown. Comparisons of the two groups were performed by Wilcoxon Rank Sum test and *P* values are shown on the right. *, *P* < 0.05 **, *P* < 0.01, N.S., no significant differences.

**Supplementary Table 2.** Amounts of bile acids in feces.

| Bile acids | Control　 (nmol / g feces) Average ± Standard deviation | PHGG　 (nmol / g feces) Average ± Standard deviation | *P* value |
| --- | --- | --- | --- |
| DCA | 81.75 ± 40.88 | 82.62 ± 15.81 | N.S. |
| LCA | 78.44 ± 22.75 | 92.66 ± 8.92 | N.S. |
| CA | 8.25 ± 5.33 | 5.24 ± 1.74 | N.S. |
| UDCA | 0.38 ± 0.25 | 0.69 ± 0.29 | * |
| a-MCA/w-MCA | 0.29 ± 0.45 | 0.19 ± 0.11 | N.S. |
| CDCA | 0.06 ± 0.14 | 0.03 ± 0.09 | N.S. |
| TCDCA | 0.04 ± 0.04 | 0.02 ± 0.03 | N.S. |
| b-MCA | 0.03 ± 0.09 | 0.00 ± 0.00 | N.S. |
| GDCA | 0.02 ± 0.03 | 0.02 ± 0.03 | N.S. |
| TLCA | 0.01 ± 0.02 | 0.03 ± 0.05 | N.S. |
| TDCA | 0.01 ± 0.03 | 0.02 ± 0.05 | N.S. |

Average values ± standard deviation for each fecal bile acid and group are shown. Comparisons of the two groups were performed by Wilcoxon Rank Sum test and *P* values are shown on the right. *, *P* < 0.05, N.S., no significant differences.

**Supplementary Table 3.** Concentration of bile acids in serum.

| Bile acids | Control (nM) Average ± Standard deviation | PHGG (nM) Average ± Standard deviation | *P* value |
| --- | --- | --- | --- |
| CA | 15220.3 ± 4298.1 | 16868.7 ± 4531.1 | N.S. |
| DCA | 1329.9 ± 219.0 | 1848.5 ± 329.4 | ** |
| GCDCA | 39.2 ± 110.9 | 157.2 ± 261.1 | N.S. |
| GDCA | N.D. | 57.8 ± 163.5 | N.S. |
| GCA | N.D. | 183.4 ± 518.8 | N.S. |

Average values ± standard deviation for each serum bile acid and group are shown. Comparisons of the two groups were performed by Wilcoxon Rank Sum test and *P* values are shown on the right. **, *P* < 0.01, N.S., no significant differences.
